# One year later: longitudinal effects of flexible school start times on teenage sleep and subjective psychological outcomes

**DOI:** 10.1101/2021.07.27.453940

**Authors:** Anna M. Biller, Carmen Molenda, Giulia Zerbini, Till Roenneberg, Eva C. Winnebeck

**Author notes:** equal contribution (Anna M. Biller, Carmen Molenda). current address: Institute of Psychology, Bundeswehr University Munich, Munich, Germany (Anna M. Biller). current address: Institute and Polyclinic for Occupational-, Social- and Environmental Medicine, Ludwig Maximilian University Munich, Munich Germany (Till Roenneberg). current address: School of Medicine, Chair of Neurogenetics, Technical University of Munich, and Institute of Neurogenomics, Helmholtz Center Munich, Munich, Germany (Eva C. Winnebeck). corresponding author, <u>Correspondence should be addressed to:</u> Eva Winnebeck, Institute of Neurogenomics, Ingolstädter Landstr. 1, 85764 Neuherberg, Germany.

## Abstract

Early school times fundamentally clash with the late sleep of teenagers. This mismatch results in chronic sleep deprivation, which poses acute and long-term health risks and impairs students’ learning. Despite conclusive evidence that delaying school times has immediate benefits for sleep, the long-term effects on sleep are unresolved due to a shortage of longitudinal data. Here, we studied whether a *flexible* school start system, with the daily choice of an 8AM or 08:50AM-start, allowed secondary school students to improve their sleep and psychological functioning in a longitudinal pre-post design over exactly 1 year. Based on 2 waves, each with 6-9 weeks of daily sleep diary, we found that students maintained their 1-hour-sleep gain on days with later starts, both longitudinally (n=28) and cross-sectionally (n=79). This sleep gain was independent of chronotype and frequency of later starts but differed between genders. Girls were more successful in keeping early sleep onsets despite later sleep offsets, whereas boys delayed their onsets and thus had reduced sleep gains after 1 year. Students also reported psychological benefits (n=93), increased sleep quality and reduced alarm-driven waking on later school days. Despite these benefits on later schooldays, overall sleep duration was not extended in the flexible system. This was likely due to the persistently low uptake of the late-start option. If uptake can be further promoted, the flexible system is an appealing alternative to a fixed delay of school starts owing to possible circadian advantages (speculatively through prevention of phase-delays) and psychological mechanisms (e.g. sense of control).

**Significance statement:** Teenage sleep becomes progressively later during adolescence but school starts do not accommodate this shifted sleep window. This mismatch results in chronic sleep deprivation in teenagers worldwide, which is a pervasive public health concern. Delaying school starts could counteract this misalignment if students keep sleep onsets stable. However, few studies have investigated long-term effects of delayed starts on sleep. We observed here that students slept persistently longer and better when school started later in a flexible start system. Girls were especially successful in keeping stable onsets. Students also benefitted psychologically and liked the flexible system despite only a modest uptake of later starts. The flexible system is an interesting alternative to a fixed delay and should receive more scientific attention.

## Introduction

Teenagers around the world are chronically sleep deprived because their late sleep timing often clashes with early school starts forcing them to get up long before their sleep has come to a natural end. Sleep is timed progressively later during adolescence because teenagers’ internal circadian phase (chronotype) markedly delays [1–3]. At the same time, sleep pressure (the homeostatic load) accumulates more slowly over the day compared to adults or younger children, making teenagers less tired in the evening [4,5]. These biological tendencies are exacerbated by non-biological factors, such as academic pressure or cultural influences to stay up late [6,7]. Evening activities then lead to longer exposure to artificial light at night which increases alertness [8–10] and further delays circadian rhythms resulting in later sleep timings. Consequently, many students do not get enough sleep during the school week and compensate their sleep loss by oversleeping on weekends. This is often accompanied by a delay of sleep timing on free days - a phenomenon called “social jetlag” [11]. Yet, even with weekend lie-ins, most teenagers do not achieve weekly sleep durations of at least 8 hours each night [12,13], the recommended minimum sleep amount at this age [14].

The consequences of short sleep are numerous biological and psychological health compromises. In the long-term, chronic sleep deprivation has been linked to metabolic, cardiovascular, and inflammatory diseases [15,16], to depressed mood and worsened emotional regulation [17–19], as well as substance use [20,21]. Social jetlag, too, is associated with metabolic syndrome, obesity and depression, as well as increased alcohol consumption and smoking [11,22–25].

The obvious solution, to simply delay school start times by a fixed amount, has gained much scientific and public attention over the past decades. Positive associations were found for sleep and sleep quality, daytime sleepiness, wellbeing and mood [26,27], concentration and attention in class, absenteeism and tardiness, and even motor vehicle accidents [28–32]. Nonetheless, policy-uptake is still rare (except for California, USA), also invoking the low evidence level of the findings and unclear long-term benefits as a reason [33,34]. Indeed, the vast majority of studies used a cross-sectional design, which does not allow to track individual changes over time and is prone to cohort effects if not randomized or very carefully adjusted [30,35]. Double-blinding, the gold standard in terms of evidence level, is, of course, inherently unfeasible in this context, and it seems almost impossible to convince schools to participate in randomization [36]. Although there are some real-life settings, such as in Uruguay or Argentina, where students are randomly assigned to morning, middle, and afternoon school shifts [37,38], this is not the case in most other countries around the world. The few longitudinal studies that exist often covered ≤6 months in their follow-ups [30] (but see [39–43]), and are thus prone to seasonal confounding. Furthermore, sleep, mood, and performance have often been assessed via one-off questionnaires, while continuous sleep recordings via daily sleep diaries and especially objective actimetry measures are scarce [30,32,41,44–47]. One notable exception is a recent study by Widome and colleagues who followed students over two years and found persisting extended sleep durations (measured with one week of actimetry) in students from schools who delayed bell times compared to students in schools which did not change [43].

We had previously investigated sleep changes and psychological benefits following a switch to a *flexible* start system – a highly overlooked start system that might offer some interesting advantages [48]. Here, we now report on the longer-term effects of this flexible system after 1 year of exposure. The flexible system was established at a German secondary school to provide flexibility on the school start time on a *daily basis*. This means that every single student in 10^th^-12^th^ grade decides *each day* if they attend the first period at 8AM or if they skip the first period and start at 08:50AM instead. In the rare case of a scheduled free second period, skipping the first period leads to a 10:15AM-start. Non-attended first periods have to be made up for within the same week during free periods or after classes. Right after the introduction of the flexible system, students had extended their sleep on days with later starts by more than 1 hour, reported better sleep quality and slightly less alarm-driven wakings [48]. Nonetheless, compared to baseline, sleep duration had not significantly increased in the flexible system overall. This was caused by students not making full use of the later start option but only choosing to start school later on a median of 2 schooldays per week, and by the fact that there were some infrequent later starts already at baseline. Here, we investigated the situation after 1 year. Did the uptake of the late option increase and thus lead to marked sleep benefits in the flexible system? Did students maintain their large sleep gains on days with later starts? Or did they adjust to the flexible system and delay their sleep times throughout the week?

## Methods and Materials

### Study Site

The study took part at the Gymnasium Alsdorf (50° 53’ N, 6° 10’ E), a secondary school in a town of ~45.000 residents in the West of Germany. A Gymnasium is the most academic of secondary schools in Germany and grants access to higher education after 8-9 years of study and successful completion of the final exam. The school received the German School Award in 2013 for its innovative teaching [49]. It follows an educational system called “Dalton plan” that incorporates daily self-study periods called “Dalton hours” during which students work through a personal 5-week curriculum with a teacher and on a subject of their own choice.

### Change in School Start Times

The school changed permanently from a fixed start (“conventional system”) to a flexible start (“flexible system”) for older students (grades 10-12) on February 1^st^, 2016. In the conventional system, the first period started at 8AM. On a median of 1 day/week, depending on students’ individual timetables, classes started with the second period at 8:50AM.

In the flexible system, one of the two daily self-study periods was advanced into the first period (lasting 08:00-08:45AM) and made optional to attend for students in grades 10-12 (for an example timetable see [48]). Students could thus choose daily whether to start at 8AM with the first self-study period or skip it and start at 08:50AM instead (called “9AM” for simplicity). On a median of 1 day/fortnight, students also had a scheduled free second period (08:50-09:50AM), i.e. the chance to turn the 08:50-start into an 10:15-start when skipping the first period (“>9AM”). Given the low frequency of 10:15-starts (median 25%, see results), we grouped the two types of later school starts into “≥9AM-days” and compared those with 8AM-days.

Students had to make up for the skipped first periods throughout the week, using gap periods or adding study time after their last classes (up to the official school closing at 4:15 PM). To be able to start later on all 5 schooldays/week, most students had to make use of both options since their individual schedules did not provide 5 gap periods and 5 early class ends per week.

### Study Design

Data were collected in two waves that were exactly one year apart (Fig. 1A). Wave 1 took place in winter 2016 and consisted of i) a baseline data collection covering 3 weeks in January (t0, Jan 8^th^ to 31^st^, 2016) in the conventional system with mainly 8AM-starts, ii) a data collection for 6 weeks (t1, Feb 1^st^ to Mar 14^th^, 2016) in the flexible system right after its introduction on Feb 1^st^, 2016. For the follow-up study (wave 2), we chose the matching photoperiod and time of t1, lasting from Feb 2^nd^ to Mar 20^th^, 2017 (t2). As the school had remained in the flexible system ever since the introduction, no second baseline just before t2 was carried out. The holiday periods over carnival between February 4^th^-9^th^, 2016 and February 23^rd^-28^th^ 2017 were excluded from the analyses.

**Figure 1.**
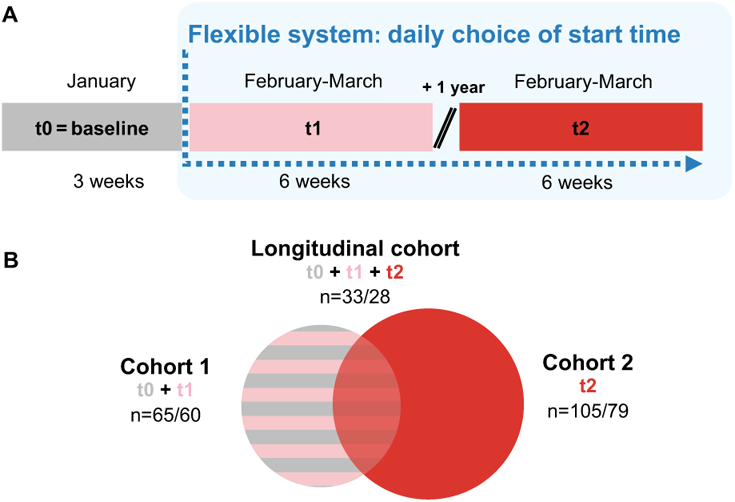
Study design and cohort overview. **A,** Schematic of longitudinal study design including wave 1 which consisted of 3 weeks of baseline assessment (t0) and 6 weeks in the flexible system (t1), and wave 2 after 1 year again covering 6 weeks of sampling (t2). **B,** Schematic of the resulting 3 different study cohorts and their respective sample sizes. Note that sample sizes vary depending on quality filters applied (see Tab.1 and participant section in methods for further information).

### Participants

Written informed consent was obtained from all participants (or their parents/guardians if <18y). The study was conducted according to the Declaration of Helsinki and approved by the school board, the parent-teacher association, the school’s student association and the ethics committee of the Medical Faculty of the LMU Munich (#774-16). We used opportunity sampling without specific exclusion criteria. In the first year (t0 and t1), 113 (45%) out of 253 possible students attending 10^th^-12^th^ grade (14-19 years) signed up, 83 (73%) students provided some data (response rate), of which 65 (70%) passed our minimal quantity and quality filter criteria (cohort 1). In the second year (t2), 162 (71%) out of 227 possible students signed up, 137 (85%) provided data (response rate), of which 105 (77%) passed the minimal filter (cohort 2). Across both years, 33 students passed the minimal filter, hence forming the longitudinal cohort. To determine the longitudinal attrition rate, one needs to note that of the 65 students in cohort 1, 16 students graduated after t1 and hence could not participate at t2 (scheduled attrition rate of 34%). Of the 49 students that could have partaken again in t2, 16 provided no or insufficient data at t2 (attrition rate of 33%). Differences in baseline characteristics between the 33 and 16 students were tested and not statistically significant (chronotype, social jetlag, gender, grade level; all p>0.05), except for age (t(47)=−2.933, p=0.005, d=0.893) with the missing students on average 0.8 years older.

Minimal filter criteria were: i) sleep information for ≥5 schooldays and ≥3 weekend days at each time point and ii) congruent, plausible data (more detailed information in [48]). For 8AM or ≥9AM-start comparisons, we additionally filtered for at least 2 8AM-days and at least 2 ≥9AM-days per person. After this additional filter, a total of 60 participants remained in cohort 1, 79 in cohort 2, and 28 in the longitudinal cohort. All students from the longitudinal cohort were granted promotion to the next grade level from wave 1 to wave 2.

### Outcome measures

#### Sleep Diary

We used a daily sleep diary (provided online via LimeSurvey.org) based on the μMCTQ [50] (a short version of the Munich Chronotype Questionnaire) and adapted it to a German student population by changing the formal you (“Sie”) to the informal you (“Du”) and work days to schooldays. Students provided sleep onset (note: not bedtime) and offset (wake time) of their past night’s sleep, whether they were woken by their alarm clock (yes/no), the type of day they woke up (schoolday or free day), when they started school (8AM, 9AM or >9AM), and their subjective sleep quality (rated on a 10-point-Likert scale from 1=“very bad” to 10=”very good”). The questionnaire did not cover any naps during the day. Although daily population of the online sleep diary was encouraged, students could also fill in data in retrospect if they had missed a day or more (they reported to have kept an offline log from which they copied their sleep timings). For more details see [48].

#### Survey

We developed a 17-item paper-pencil survey about the flexible system, which was distributed at the end of t2 and filled out by ~90% of cohort 2. Because some students did not answer all questions on the survey, the sample size ranged from 91 to 93 depending on the item. The first 7 items of the survey asked whether i) students were satisfied with the flexible system (yes/no), ii) they would rather have the old system with fixed school starts back (yes/no), iii) it was difficult for them to go to school at 8AM (never/most of the time/always), iv) it was easier to go to school at 9AM compared to 8AM (never/most of the time /always), v) how often (0 days/1-2 days/3-4 days/5 days) and vi) on which days of the week they attended the first period at 8AM (Mo/Tu/We/Th/Fr), and vii) reasons for starting school at 8AM. Answer options for vii) were to mark at least one of nine alternatives (easier to study/easier to get to school/additional study time/friends/specific teacher/specific subject/fulfill self-study quota/parents/late school end) and/or to name other reasons.

The last 10 items asked for ratings on 8AM versus ≥9AM-days. Questions were about i) sleep duration (h), ii) sleep quality (1=bad, 5=good), iii) number of schooldays with alarm-driven waking (0-5 days), iv) how tired the students felt (1=not at all, 5=very), v) ability to concentrate in class (1=bad, 5=good), vi) ability to study at home after school (1=bad, 5=good), vii) motivation to actively take part in class (1=not at all, 5=very), viii) how well they remembered new class content (1=not at all, 5=very), and ix) attitude towards school (1=negative, 5=positive). Items ii) and iv)-ix) were scored on a Five-point Likert scale.

### Data Analysis

Analyses were performed in SPSS Statistics (IBM, versions 24 and 25), R (versions 3.6.1 and 3.6.3) and R studio (versions 1.1.463, 1.2.1335 and 1.2.5042). Graphs were produced using Graph Pad Prism (versions 6 and 7) and the R package *ggplot2* [51]. Main figures (except Fig. 3) show results from the longitudinal cohort (n=28-33); results from cohort 2 (n=79-105) are provided in the text and SI.

**Figure 3.**
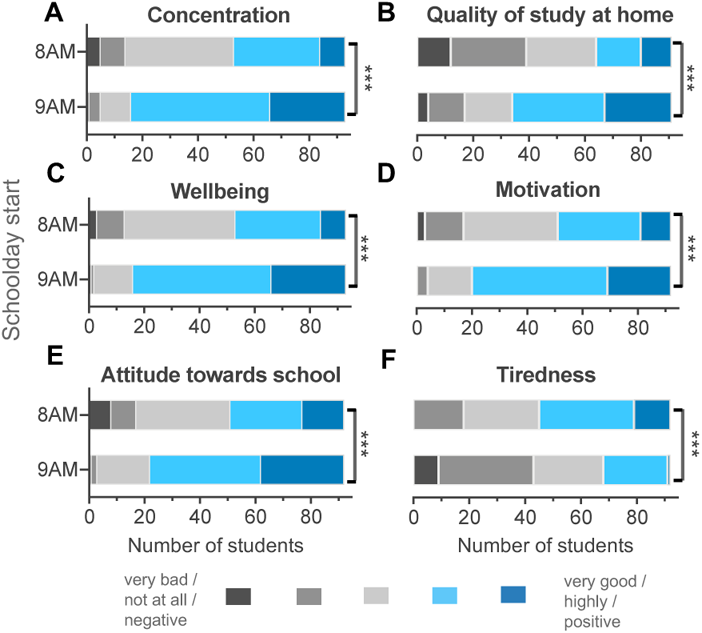
Comparison of subjective psychological benefits between 8AM-days and ≥9AM-days in the flexible system. Results from the survey at end of wave 2 asking cohort 2 for the following ratings: **A,** ability to concentrate during class (Z=6.419, d=1.784, n=93), **B,** quality of study at home after school (Z=6.055, d=1.643, n=91), **C,** general wellbeing (Z=6.559, d=1.855, n=93), **D,** motivation to attend school (Z=5.927, d=1.572, n=92), **E,** attitude towards school (Z=5.896, d=1.545, n=92), and **F,** tiredness during class (Z=5.419,d=1.369, n=92). Wilcoxon signed rank test, *p<0.05, **p<0.01, ***p<0.001.

#### Sleep Data

Daily sleep data from diaries were aggregated as mean per person for 10 conditions: at t0 for schooldays and weekends; and at t1 and t2 for schooldays, weekends, 8AM-days, and ≥9AM-days. From these aggregates, we derived the following variables as per equations below for each of the time points (t0, t1, t2): average daily sleep duration during the week (SD_week_), chronotype as midsleep on free days (MSF) corrected for oversleep (MSF_sc_), and social jetlag (SJL); for t1 and t2 only: absolute difference between ≥9AM-days and 8AM-days for variables of interest (DELTA x), frequency of alarm-driven waking, and frequency of ≥9AM-starts.

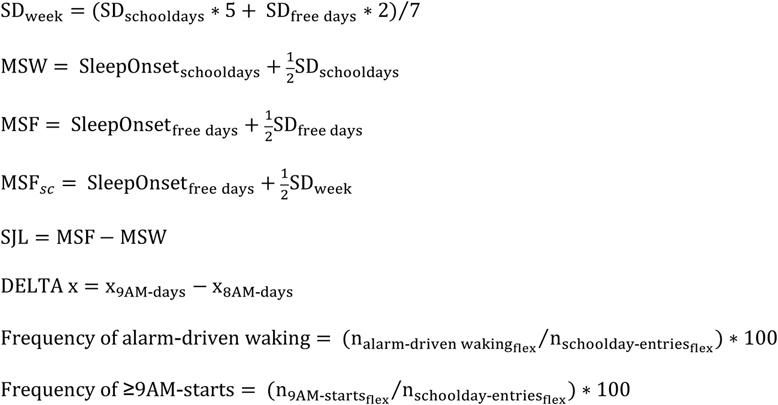

#### Statistical analyses

Unless indicated otherwise, descriptive statistics are reported as mean ± standard deviation and test statistics, p-values and effect size measures are abbreviated as follows: *t*, t-test; *Z*, Wilcoxon signed-rank test; *F*, ANOVA; *r*, Pearson correlation; *rho*, Spearman rank correlation; *p*, significance level; *d*, Cohen’s d; *d_z_,* Cohen’s d for paired t-tests; *b*, unstandardized coefficient; *beta*, standardised coefficient. Significance levels were set to *p*<0.05 for all statistical analyses. All data were tested on normality (histograms, QQ plots, Shapiro-Wilk’s test) and sphericity (Mauchley’s test; Greenhouse-Geisser corrections used for ANOVA if violated). If normality was violated, non-parametric tests were performed except for ANOVA analysis since violations were marginal.

Group difference for attrition groups were tested via independent t-test (chronotype, social jetlag, age) or Chi squared test (gender, class).

For sleep variables in the longitudinal cohort, we performed 1-way repeated measures ANOVAs with the factor time point (t0/t1/t2), 2-way repeated measures ANOVAs with the factors time point (t1/t2) and school start (8AM/≥9AM-days), and with the factors time point (t0/t1/t2) and day (schooldays/weekend). For sleep variables in cohort 2, paired t-tests (two-sided) were run for school start (8AM/≥9AM-days) and days (schooldays/weekend), and Wilcoxon signed rank test for sleep quality and survey items. Gender differences in sleep variables were assessed via 2-way mixed ANOVA with gender (female/male) and time point (t1/t2), and via linear regression (including the covariates grade level, chronotype and frequency of ≥9AM-starts) for DELTA sleep duration/onset/offset using the *nlme* package in R [52]. ANOVA results are presented above each graph (main effects and interaction). If the main interaction was significant, we interpreted (and thus provide) only the simple effects instead of the main effects. In cases of three levels within one factor, necessary post hoc tests were carried out using Bonferroni corrections.

Pearson and Spearman rank correlations were performed to assess associations between DELTA sleep duration and chronotype or frequency of ≥9AM-starts, respectively. Frequency of alarm driven waking was analysed using logistic regression (*lme4* package R [53]). Due to a large ceiling effect, we dichotomised this variable based on a median split at 100%-use (<100%: “less use”) and accommodated the repeated measures nature of the data by including ID as a random effect in a mixed regression model. Gender was included as covariate (males were woken more often by an alarm than females in the flexible system) but gender did not reach statistical significance.

## Results

During the first wave of our study [48], we had monitored students’ sleep in detail via diaries and actimetry for 3 weeks during baseline (=t0) and 6 weeks immediately after the change into the flexible system (=t1). To investigate the longer-term effects, we conducted the second wave after exactly 1 year (t2) at the same photoperiod as t1 to optimally control for seasonal effects. After 6 weeks of daily sleep diary, we also surveyed subjective wellbeing and psychological functioning on days with early versus later starts.

We allowed students to take part in wave 2 (Fig. 1A) irrespective of their participation beforehand, so our study eventually consisted of three cohorts (Fig. 1B): (i) cohort 1 provided sleep data at t0 and t1 (n=60-65), (ii) cohort 2 provided sleep and survey data only at t2 (n=79-105), and (iii) the longitudinal cohort provided sleep data throughout from t0-t2 (n=28-33; Tab. 1 and Fig. 1B). The samples sizes within each cohort varied due to different filters employed for different analysis questions (see participant section in methods).

**Table 1.** Composition of study cohorts. Displayed are cohort characteristics after standard filter criteria. An additional filter (see participant section in methods) was applied for comparisons between 8AM and ≥9AM-days, which reduced cohort 1 to 60 students, the longitudinal to 28 students, and cohort 2 to 79 students. Abbreviations: n, number of individuals; SD, standard deviation; IQR, interquartile range; conv., conventional.

|  |  | Cohort 1 | Longitudinal cohort |  | Cohort 2 |
| --- | --- | --- | --- | --- | --- |
| Time points |  | t0 and t1 | t0 and t1 | t2 | t2 |
| Participants |  |  |  |  |  |
| Total | n | 65 | 33 |  | 105 |
| Females | n (%) | 40 (62%) | 20 (60%) |  | 73 (70%) |
| Grade level | n (%) per level<br>10<sup>th</sup>/11<sup>th</sup>/12<sup>th</sup> | 26/23/16<br>(40/35/25%) | 20/13/0<br>(60/40/0%) | 0/20/13<br>(0/60/40%) | 29/38/38<br>(28/36/36%) |
| Age | mean<br>(SD, range) | 16.5<br>(1.2, 14–19) | 15.8<br>(0.9, 14-17) | 16.9<br>(0.9, 15-18) | 16.7<br>(1.1, 15-21) |
| Chronotype<br>(MSF<sub>sc</sub>; time in h) | mean<br>(SD, range) | 4.6<br>(0.9, 2.1–7.0) | 4.3<br>(0.7, 2.1-5.9) | 4.6<br>(0.9, 0.8-6.2) | 4.7<br>(1.0, 0.2-8.6) |
| Social jetlag (h) | mean<br>(SD, range) | 1.8<br>(0.7, 0.3-3.8) | 1.7<br>(0.6, 0.3-3.1) | 1.9<br>(0.6, 0.5-3.3) | 2.0<br>(0.8, 0.2-6.0) |
| Sleep duration (h) | mean<br>(SD, range) | 7.6<br>(0.8, 5.2-8.9) | 7.7<br>(0.8, 5.2-8.8) | 7.6<br>(0.7, 6.1-9.0) | 7.7<br>(0.7, 6.1-9.3) |
| Proportion of schooldays with later starts |  |  |  |  |  |
| $\geq 9$AM-use | median<br>(IQR) | 32%<br>(19-55) | 39%<br>(20-51) | 22%<br>(11-46) | 24%<br>(10-47) |
| Students reaching $\geq 8$ hours of sleep in the flexible system | | | | | |
| 8AM-days | % | 15.4% | 12.0% | 3.0% | 6.8 % |
| $\geq 9$AM-days | % | 50.0% | 59.4% | 45.2% | 47.3% |
| Schooldays | % | 18.5% | 18.2% | 9.1% | 13.3% |
| Weekends | % | 73.8% | 84.8% | 69.7% | 74.3% |
| Number of students per outcome |  |  |  |  |  |
| Sleep | 8AM vs. $\geq 9$AM-days | n=60 | n=28 | | n=79 |
|  | conv. vs. flexible system | n=65 | n=33 |  | n=105 |
| Psychological benefits |  |  |  |  | n=91-93 |

### Frequency of later starts (≥9AM-use)

Notably, our participants accumulated fewer late starts per week than expected. We had observed this for cohort 1 [48] but now saw this confirmed in cohort 2, where participants (n=105) chose to skip the first period only on a median of 24% of their schooldays (IQR: 10-47), which equates to 1.2 ≥9AM-days per 5-day school week. Similarly, the longitudinal cohort (n=33) had a median frequency of late starts (“≥9AM-use”) of 39% (20-51) and 22% (11-46) during t1 and t2, with no systematic difference between the time points (Z=−1.653, p=0.098). Importantly, ≥9AM-use varied drastically between individual participants from 0% to 100% of their schooldays, with 8:50AM-starts making up the majority of later starts per person and 10:15AM-starts, due to a second free period, only 25% (median, IQR: 6.3-60).

#### Sleep on days with later school starts

In the following, we present analyses *within* the flexible system comparing days with early school starts (“8AM-days”) to those with later starts (“≥9AM-days”).

#### Student slept longer and better on days with later school starts – an improvement persisting over one year

How was students’ sleep altered by later school start times in the flexible system over one year? We showed previously that, right after the introduction of the flexible system, students from cohort 1 slept about one hour longer on ≥9AM-days by maintaining their sleep onset but delaying their sleep offset [48]. After one year, we found the same sleep gain of ~1h for cohort 2 and, importantly, also in the longitudinal cohort across both time points. Repeated measures ANOVAs in the longitudinal cohort (n=28) showed that sleep onsets did not differ with school start time or time point (Fig. 2A), whereas sleep offsets were 61 min (± 47) later on average (Fig. 2B), and students hence slept 62 min (± 47) longer on ≥9AM-days compared to 8AM-days across both time points (Fig. 2C and 2F, full statistics in Figures). Findings from cohort 2 (n=79) tally with this pattern: sleep onsets on 8AM and ≥9AM-days were comparable (t[78]=−1.87, p=0.065; d_z_=0.210), while wake up times were significantly later on ≥9AM-days (t[78]=−19.75, p<0.001, d_z_=2.222), which resulted in 60 min longer sleep durations on those days (t[78]=−10.83, p<0.001, d_z_=−1.218). This large sleep gain likely results from ≥9AM-days incorporating not only 8:50-starts (75%) but also some 10:15-starts (25%).

**Figure 2.**
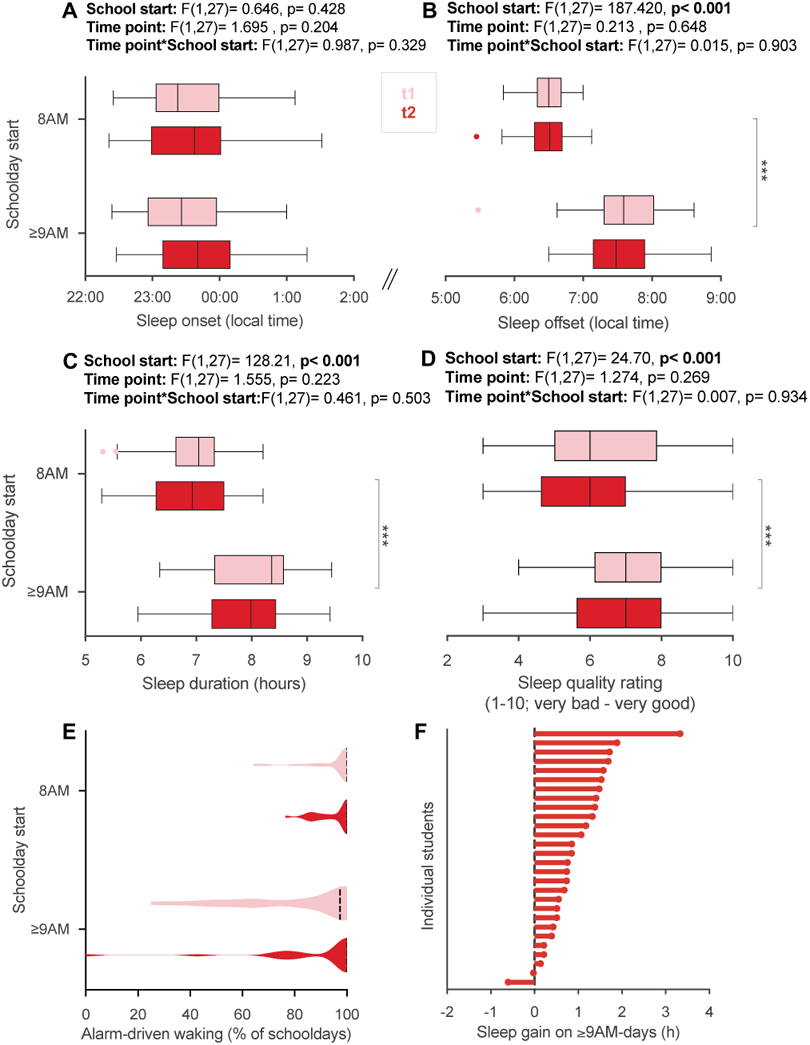
Comparison of sleep parameters between 8AM-days and ≥9AM-days in the flexible system. **A-F,** Sleep parameters from the longitudinal cohort (n=28) comparing 8AM and ≥9AM-days at t1 (light red) and t2 (dark red) intra-individually. **A,** Average sleep onset, **B,** offset, **C,** duration, and **D,** quality on 8AM versus ≥9AM-days in the flexible system across time points. Results of two-way repeated measures ANOVA with the within-subject factors school start (8AM/≥9AM) and time point (t1/t2) are reported above each graph. Brackets indicate statistically significant post-hoc comparisons. **E,** Proportion of schooldays with alarm-driven waking. **F,** Sleep gain on ≥9AM-days at t2 for each student. Depicted is the average absolute difference in sleep duration between 8AM and ≥9AM-days. Positive values mean longer sleep on ≥9AM-days. Dashed lines in violin plots show medians. All boxplots are Tukey boxplots. *p<0.05, **p<0.01, ***p<0.001.

Furthermore, subjective sleep quality was improved on ≥9AM-days by 1 point on a 10-point Likert scale for cohort 1 [48] and cohort 2 (n=79, Z=−5.874, p<0.001, d=−1.761), and also longitudinally across time points (n=28, Fig. 2D). In addition, the extensive use of alarm clocks remained slightly reduced on ≥9AM-days also one year into the system (Fig. 2E). Just as in cohort 1 [48], the odds for less alarm-driven waking were increased in cohort 2 (n=79, OR = 1.9, 95% CI = 1.3-4.1) and showed a similar qualitative pattern also in the longitudinal cohort (n=28; Fig. 2E), demonstrating that a natural waking was more likely when school started later.

#### Students reported profound improvements in cognitive and psychological parameters on days with later school starts

To assess psychological benefits, we used survey data from the end of t2, which were provided by 90% of cohort 2. Students’ subjective ratings of their sleep, cognition and well-being on 8AM-days compared to ≥9AM-days showed statistically significant improvements in all areas assessed (n=91-93; full statistics in Fig. 3). On days with later starts, students felt generally better, less tired during class, more motivated to actively take part in class, and were better able to concentrate. Students also reported a more positive attitude towards attending school and higher quality of self-study after school.

#### Girls maintained their sleep benefit from later school starts more than boys after one year in the flexible system

We wondered whether particular students benefitted more or less than others from later starts. Therefore, we assessed the relationship of chronotype, ≥9AM-use and gender with the core sleep benefit, the sleep gain on ≥9AM-days (the difference in sleep duration between ≥9AM- and 8AM-days). In the longitudinal cohort (n=28; see further below for cohort 2), 93% of students experienced a sleep gain on ≥9AM-days across both time points (Fig. 2F), so the sleep benefit was close to universal. Chronotype was not correlated with sleep gain (t1: r=−0.024, p=0.903; t2: r=−0.091, p=0.647), i.e. both early and late chronotypes appear to have benefitted equally from later starts (Fig. 4A-B). We had already observed this in cohort 1 [48] and interpreted it as the consequence of the severe sleep deprivation in adolescent students which afflicts even earlier chronotypes. Similarly, no matter how often the students attended school later, their sleep gain on ≥9AM-days seemed not systematically affected. Although correlations indicated smaller gains with more frequent ≥9AM-use at t1 (rho=−0.55, p=0.003), this was mainly driven by two over-benefitting individuals with low ≥9AM-use and one under-benefitting individual with high use – all three identified as outliers already in our wave-1-analyses [48]. Without these three Tukey outliers, the relationship was smaller and statistically non-significant (rho=−0.37, p=0.064). There was also no correlation between sleep gain and ≥9AM-use during t2 (rho=0.028, p=0.889; Fig. 4A-B).

**Figure 4.**
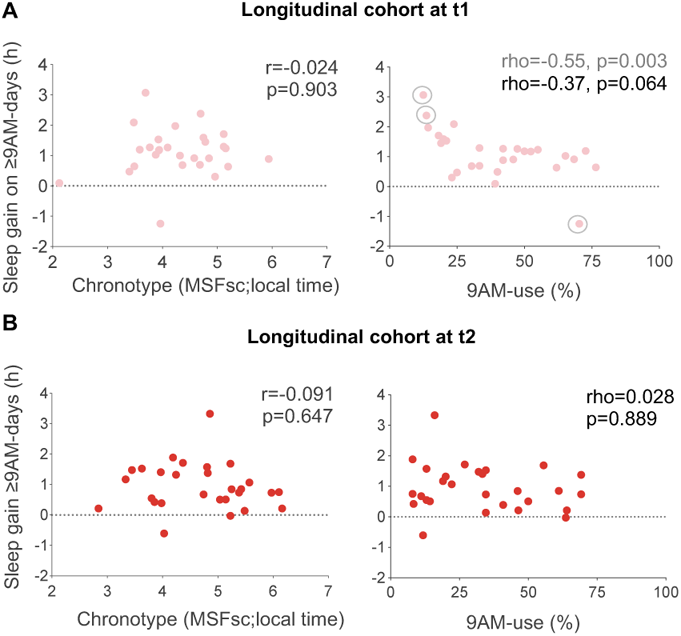
Inter-individual differences in sleep gain on ≥9AM-days. Shown are relationships between chronotype (MSFsc; local time) or frequency of ≥9AM-starts (% of schooldays with later starts) with sleep gain on ≥9AM-days. Sleep gain was quantified as the absolute difference in sleep duration between ≥9AM and 8AM-days, with positive numbers indicating longer sleep duration on ≥9AM-days. Data are from the longitudinal cohort (n=28) during **A,** t1 (light red) and **B,** t2 (red). Results of Pearson (r) and Spearman (rho) correlations are indicated. Tukey outliers in sleep gain, which overproportionally influence the correlation with 9AM-use at t1 (right panel in A), are marked with grey empty circles, and correlation results including (grey) and excluding outliers (black) are provided.

In contrast, gender showed a clear effect on sleep gain after one year: both genders enjoyed similar sleep gains during t1, as also found in cohort 1 [48], but boys clearly reduced their sleep gain during t2 from 1.3h (± 0.53) to 0.5h (± 0.53, detailed statistics in Fig. 5C and Tab. S1). Follow-up analyses revealed that the reduced sleep gain in boys resulted from a delay in their sleep onsets on ≥9AM-days compared to 8AM-days (Fig. 5A), while their offset times were unaltered during t2 (n=28; Fig. 5B, statistics in Fig. 5A and Tab. S1).

**Figure 5.**
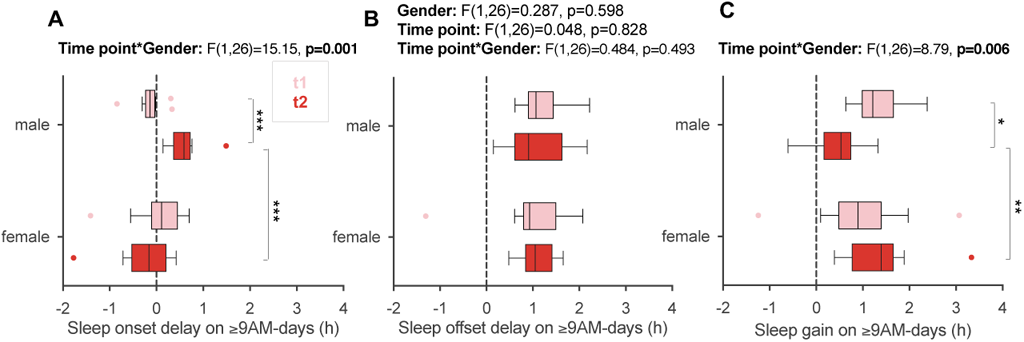
Gender differences in sleep onset, offset and duration on ≥9AM-days versus 8AM-days in the flexible system. Depicted is the average absolute difference between 8AM and ≥9AM-days in **A,** sleep onset (sleep onset delay) and **B,** sleep offset (sleep offset delay) and **C,** sleep duration (sleep gain) for the longitudinal cohort (n=28), with positive numbers indicating higher values on ≥9AM-days. Results of two-way mixed ANOVAs with the between-subjects factor gender (female/male) and the within-subjects factor time point (light red=t1/red= t2) are reported above each graph. Given the significant interaction effect on sleep onset delay, main effects are not reported, instead statistically significant post-hoc comparisons are indicated. See main text and SI for detailed effect sizes. All boxplots are Tukey boxplots. *, p<0.05; **, p<0.01; ***, p<0.001

The bigger sample size of cohort 2 (n=79) allowed us to address all the above relationships together in single regression models, in particular the reasons for the gender disparity. Besides gender, chronotype and ≥9AM-use, we also included grade level (inherently incorporating age) as predictors for sleep gain, sleep onset delay and sleep offset delay (the differences between ≥9AM and 8AM-days; Tab. S2). The regression results corroborated all observations from the longitudinal cohort showing that only gender had a significant influence on any of the outcomes, namely sleep gain and sleep onset delay (Tab. S2). Boys reduced their sleep gain on average by 0.52 h (b=−0.52, p=0.010, r=>0.6), which was driven by a delay in their onset on ≥9AM-days by 0.53h (b=0.53, p<0.001, r>0.6), while their offset was unchanged (b=0.01, p=0.942, r=0.07). Sensitivity analyses indicated that this effect was not just driven by the longitudinal cohort comprising 35% of cohort 2. Taken together, while most inter-individual differences did not systematically influence sleep gains, boys showed a delay in sleep onset and thus displayed a smaller sleep gain on ≥9AM-days after one year in the flexible system.

### Sleep in the flexible system versus baseline

Despite obvious improvements in sleep and subjective parameters on ≥9AM-days also after one year, it is essential to determine if these actually translated into better sleep in the flexible system overall. Based on our analyses of cohort 1 [48], this was largely not the case during the first six weeks after the introduction of the flexible system. Most likely, the limited ≥9AM-use in combination with occasional late starts during baseline reduced improvements by the flexible system compared to the conventional system. But did long-term effects emerge after one year of exposure in the flexible system?

#### Students did not extend their sleep in the flexible system overall

Analyses in the longitudinal cohort (n=33) revealed that students’ sleep was not improved compared to baseline even after 1 year in the flexible system. Despite small delays in sleep offset on schooldays (Fig. 6A, detailed statistics in Fig. 6 and Tab. S3), sleep duration on schooldays and across the week were not significantly increased at t1 or t2 compared to t0 (Fig. 6B). Students still only slept 7.6 h (± 0.65) on a daily average across the week (including weekend catch-up sleep) at t2, a sleep duration below the recommended 8-10 h for this age group [54]. Students’ chronotype remained expectedly late across all time points (Fig. 6C), and there was still a substantial difference between sleep timing on schooldays and weekends (Fig. 6A; Tab. S4 for similar results in cohort 2). Students’ social jetlag, which quantifies this typical shifting between the ‘schoolday-time zone’ and the ‘weekend-time zone’, although reduced at t1 by 30 min (± 0.62, p=0.002), was indistinguishable from baseline after one year (p=0.256; Fig. 6D). So, the mild reduction in social jetlag experienced immediately after entering the system was lost later on, emphasizing that there was no widespread improvement in sleep under the low ≥9AM-use in the flexible system.

**Figure 6.**
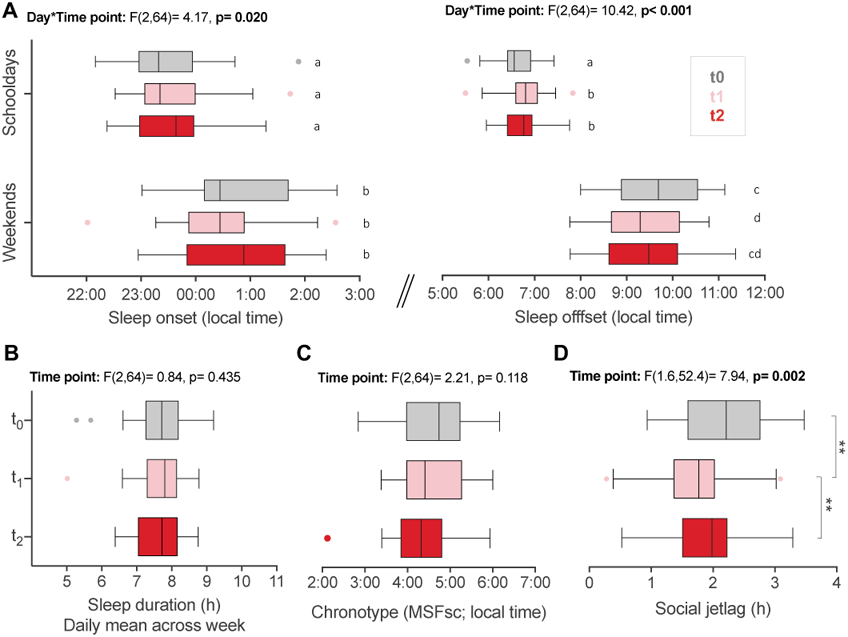
Comparison of sleep parameters across school start systems. Sleep parameters from the longitudinal cohort (n=33) comparing the conventional start system at baseline (t0, grey) with the flexible system during t1 (light red) and t2 (dark red). **A,** Average sleep onset and offset on schooldays and weekends. Results of two-way repeated measures ANOVAs with the factors day (schooldays/weekends) and time point (t0/t1/t2) are provided. Given the significant interaction effect, main effects are not reported. Letters indicate results of post-hoc tests on simple contrasts, with data marked by different letters demonstrating significant differences. **B,** Average daily sleep duration across the week (weighted for 5 schooldays and 2 weekend days), **C,** average chronotype, **D,** average social jetlag. Results of one-way repeated measures ANOVAs across time points are presented above each graph. Brackets indicate statistically significant post-hoc comparisons. All boxplots are Tukey boxplots. See main text and Tab. S3 for detailed effect sizes. *p<0.05, **p<0.01, ***p<0.001.

## Discussion

Teenagers show restricted sleep on schooldays and catch-up-sleep on weekends. Early school starts are a major determinant of this pattern, thereby impacting students’ daily lives and their future trajectories. Most studies that looked at delayed school starts and sleep improvements were cross-sectional (and thus could not track individual differences over time) and analysed outright and fixed delays in start times. Here, we investigated whether a flexible school start system allowed teenagers to reduce their sleep deprivation long-term, and whether this system was associated also with subjective improvements in psychological parameters.

The few studies that recorded sleep changes longitudinally after a delay in school start times reported mixed results. Bowers and Moyer (2016) determined in a meta-analysis [30] that all five longitudinal studies examined showed sleep extensions after a school start delay, and this benefit persisted until the follow-up period at 0.25 to 6 months after the delay [31,44,47,55,56]. Lo et al. (2018) also tracked sleep after a 45-min delay and found a delay in bedtime of 23 min which was sustained after 9 months [41]. In contrast, Thacher and Onyper (2016) showed a 20-min sleep extension after 45-min delay disappeared after 1 year because students delayed their sleep times [40]. Das-Friebel et al. (2020) also provided evidence that students merely shifted their sleep timing to later and thus did not benefit from their 20-min school delay after 1 year [42].

Here, in the flexible start system compared to the conventional start system, we found no shift in sleep timing but also no net sleep gains, which is probably connected to the low uptake of later starts of only 1-2 days per week on average and to occasional later starts already occurring during the conventional system. We had identified three main reasons for this low uptake via survey answers during wave 1: students could not fulfil their quota of 10 self-study periods per week without otherwise getting home later in the afternoon (75%), it was easier to get to school for the 8AM-start (40%), and students wanted to have more time to study (27%) [48]. During wave 2, these reasons remained the most common ones (54%, 37%, 50% respectively) for going early - although yet another year later the uptake of the late-start option apparently rose to a median of 79% (IQR=70-86), i.e. 4 days per week, according to data provided by the school. It is therefore likely that the temperate use of the flexible starts during our recording period underlies the persistent absence of sleep benefits in the flexible system in our sample. Thus, more late starts are probably required to translate into net sleep benefits in a flexible system. Alternatively – or in addition – the flexible system might have compensated a potential deterioration in sleep with increasing age or adolescence [57,58] and the absence of a net change in sleep between all time points is actually a success as it prevented a worsening. Longitudinal observational data, however, are unfortunately not suited to answer this question.

Within the flexible system, our results demonstrate that sleep length on ≥9AM-days remained increased on average by 1 hour even after one year, and that ≥9AM-starts were subjectively helpful for students across many psychological domains. The sleep and psychological effects might be either downstream of each other (e.g. longer and better sleep improving well-being and concentration or vice versa) or parallel improvements (e.g. more self-determination in the flexible system improving both sleep and psychological aspects in day-time functioning). The finding that almost every single student profited from a later start highlights the pervasiveness and severity of sleep deprivation in this age group.

Importantly, however, while girls’ sleep benefit on ≥9AM-days was completely sustained over the follow-up period, boys’ sleep gain was reduced after 1 year since they fell asleep later on ≥9AM days than on 8AM-days at t2. This could have been a cohort effect of the small longitudinal cohort but the larger cohort 2, which had a similar gender ratio, showed the same pattern. The delay in sleep onsets for boys but not girls is a central finding, since avoiding delays in sleep onsets is key to long-term success of later school start times, both flexible and fixed. Our analyses revealed no effects of chronotype or frequency of later starts on this delay. We can thus only speculate about the possible biological, psychological and behavioural reasons explaining the observed gender difference, ranging from different circadian light sensitivities [59] to (un)consciously differing sleep hygiene or pre-bed activities (e.g. bed procrastination that has been shown to be higher in males [60]). It is clear that this gender difference after 1 year raises many central questions and might underlie the contradictory findings from the few previous longitudinal studies (with e.g. all-girls samples [41] or few gender analyses), highlighting the urgent need for long-term follow-ups of sleep timing adjustments with differential effects.

The benefits of later school starts are also reflected by the fact that 45% to 59% of students across all cohorts enjoyed at least 8 hours of sleep on ≥9AM-days (Tab. 1), while numbers looked worrying on 8AM-days, when only 3% to 15% of students reached the minimal amount of 8h required for healthy sleep in teenagers [54]. Although students still did not get the recommended 8-10h on schooldays overall, this demonstrated that later starts are beneficial for teenage sleep and constitute a move in the right direction. Sleep lengths on ≥9AM-days got closer to more optimal levels, which we otherwise only observed on weekends when 70-85% of students in our sample reached at least 8h of sleep. Other studies found similar effects only when school started much later, such as in the afternoon [61,62].

Another bonus is that students themselves liked the new system. They were more motivated to go to school, they rated their concentration and motivation higher during class, and generally felt better on ≥9AM-days. These are also prerequisites for good academic learning and achievement. Whether the flexible system was associated with an improvement in students’ grades was analysed in detail in an accompanying manuscript [63].

Our study has some limitations that have not yet been mentioned. Sleep analyses were solely based on subjective diaries entries. However, importantly, diary data corresponded very well to objective activity data in cohort 1 (r=0.8-0.9) [48], and other studies report similar correlations [64,65], so we assume faithful reporting from our sample. Furthermore, our sleep calculations did not consider potential naps and hence might underestimate the total sleep duration in some students. Finally, we also did not have data on the socioeconomic background of our participants. Students attending Gymnasium (the most academic type of school in Germany) tend to be from families with higher socio-economic status, and often at least one parent has a similar educational level (65.9% of parents obtained A-levels, and 22.2% a General Certificate of Secondary Education equivalent [66]).

In conclusion, we showed that students maintained a 1-hour sleep gain on days with later starts over a period of one year in a flexible system with clear subjective psychological benefits. Flexible school starting times could therefore become an interesting alternative to later starts to support sleep and mental and physical health of secondary school students. Additionally, teaching students to take responsibility, which incorporates to decide for themselves when to learn and to some extent when to start school, could increase their motivation, investment, and wellbeing, and can thus have potential indirect effects on their sleep quality.

## Acknowledgements

We thank the students for their participation and the management of the Gymnasium Alsdorf, in particular W Bock and O Vollert, for their open support throughout the study. We also thank Dr. M Vuori-Brodowski for her contributions during wave 1, general advice and statistical support.

## Author contributions (CRedIT Taxonomy)

Conceptualisation: TR, ECW Methodology: ECW, AMB, CM Investigation: AMB, CM, ECW Data curation: AMB, CM

Formal analysis: AMB, CM, ECW Validation: ECW, AMB, CM, GZ Supervision: ECW

Visualisation: AMB, CM, ECW Writing – original draft: AMB, ECW

Writing – review and editing: AMB, ECW, CM, GZ

## Data availability

Data were collected with a consent form that prohibits online deposition of data for open access sharing. This prohibition was implemented in order to protect participants’ privacy in a cohort where most individuals are well-acquainted with each other and peers or teachers might identify participants. Data are available from the corresponding author upon reasonable request.

## Supplementary information

**Table S1.**
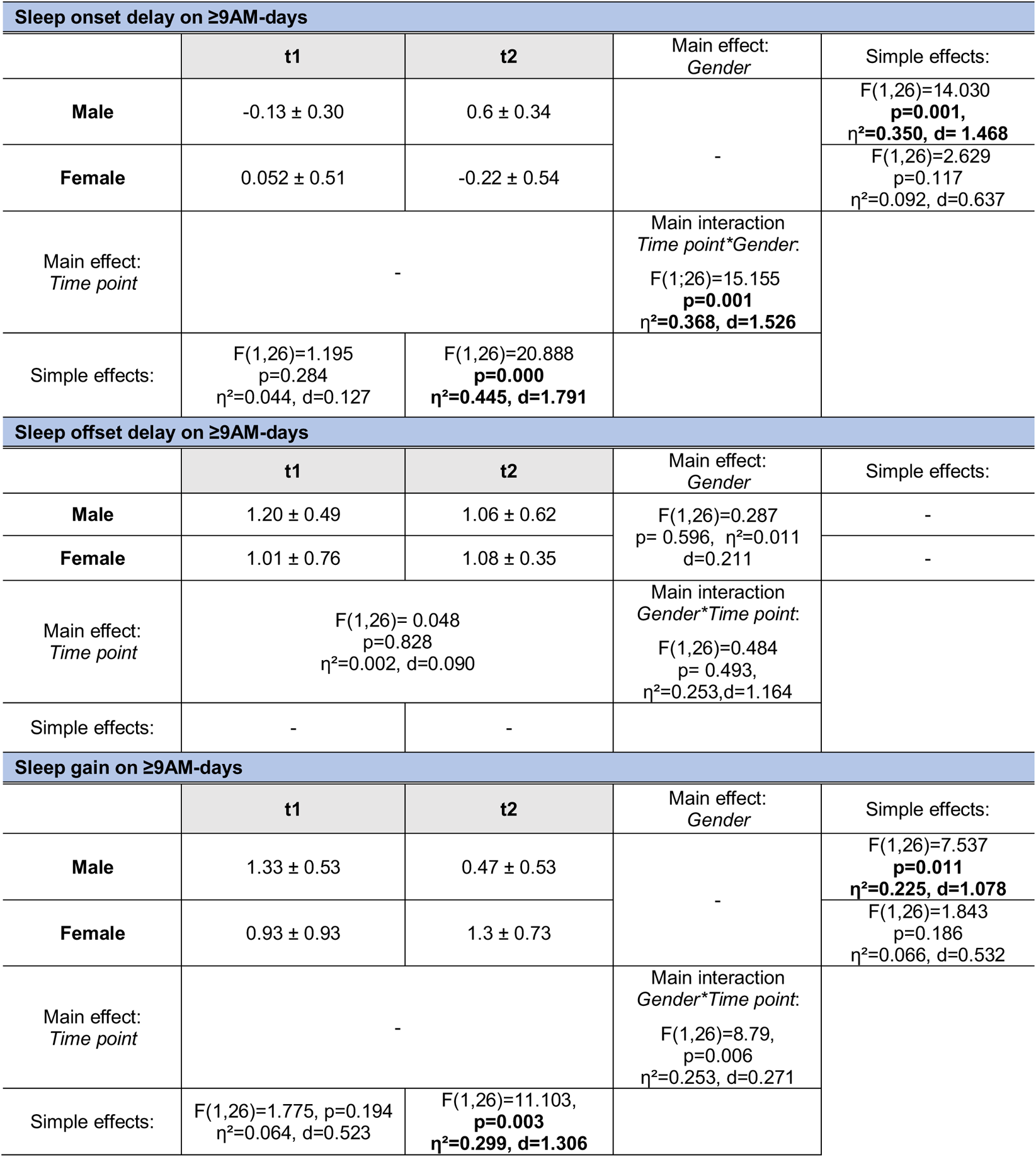
Sleep differences between time points and gender in the longitudinal cohort (post-hoc comparisons relating to Fig. 5). Two-way mixed ANOVAs were run for sleep onset delay, sleep offset delay, and sleep gain on ≥9AM-days respectively with the within-factor time point (t1/t2) and between-factor gender (male/female). In case of significant interaction of both factors, simple effects were indicated instead of interpreting the main effects. Data presented are mean ± standard deviation from the longitudinal cohort (n=28). η^2^, (partial) eta squared; d, Cohen’s d.

| Sleep onset delay on $\geq 9$AM-days | | | | |
| --- | --- | --- | --- | --- |
|  | t1 | t2 | Main effect:<br>*Gender* | Simple effects: |
| Male | -0.13 ± 0.30 | 0.6 ± 0.34 | - | F(1,26)=14.030<br>**p=0.001**,<br>$\eta^2=0.350$, d= 1.468 |
| Female | 0.052 ± 0.51 | -0.22 ± 0.54 | | F(1,26)=2.629<br>p=0.117<br>$\eta^2=0.092$, d=0.637 |
| Main effect:<br>*Time point* | - | | Main interaction<br>*Time point\*Gender*:<br>F(1;26)=15.155<br>**p=0.001**<br>$\eta^2=0.368$, d=1.526 | |
| Simple effects: | F(1,26)=1.195<br>p=0.284<br>$\eta^2=0.044$, d=0.127 | F(1,26)=20.888<br>**p=0.000**<br>$\eta^2=0.445$, d=1.791 | | |
| Sleep offset delay on $\geq 9$AM-days | | | | |
|  | t1 | t2 | Main effect:<br>*Gender* | Simple effects: |
| Male | 1.20 ± 0.49 | 1.06 ± 0.62 | F(1,26)=0.287<br>p= 0.596, $\eta^2=0.011$<br>d=0.211 | - |
| Female | 1.01 ± 0.76 | 1.08 ± 0.35 | | - |
| Main effect:<br>*Time point* | F(1,26)= 0.048<br>p=0.828<br>$\eta^2=0.002$, d=0.090 | | Main interaction<br>*Gender\*Time point*:<br>F(1,26)=0.484<br>p= 0.493,<br>$\eta^2=0.253$,d=1.164 | |
| Simple effects: | - | - |  |  |
| Sleep gain on $\geq 9$AM-days | | | | |
|  | t1 | t2 | Main effect:<br>*Gender* | Simple effects: |
| Male | 1.33 ± 0.53 | 0.47 ± 0.53 | - | F(1,26)=7.537<br>**p=0.011**<br>$\eta^2=0.225$, d=1.078 |
| Female | 0.93 ± 0.93 | 1.3 ± 0.73 | | F(1,26)=1.843<br>p=0.186<br>$\eta^2=0.066$, d=0.532 |
| Main effect:<br>*Time point* | - | | Main interaction<br>*Gender\*Time point*:<br>F(1,26)=8.79,<br>p=0.006<br>$\eta^2=0.253$, d=0.271 | |
| Simple effects: | F(1,26)=1.775, p=0.194<br>$\eta^2=0.064$, d=0.523 | F(1,26)=11.103,<br>**p=0.003**<br>$\eta^2=0.299$, d=1.306 | | |

**Table S2.**
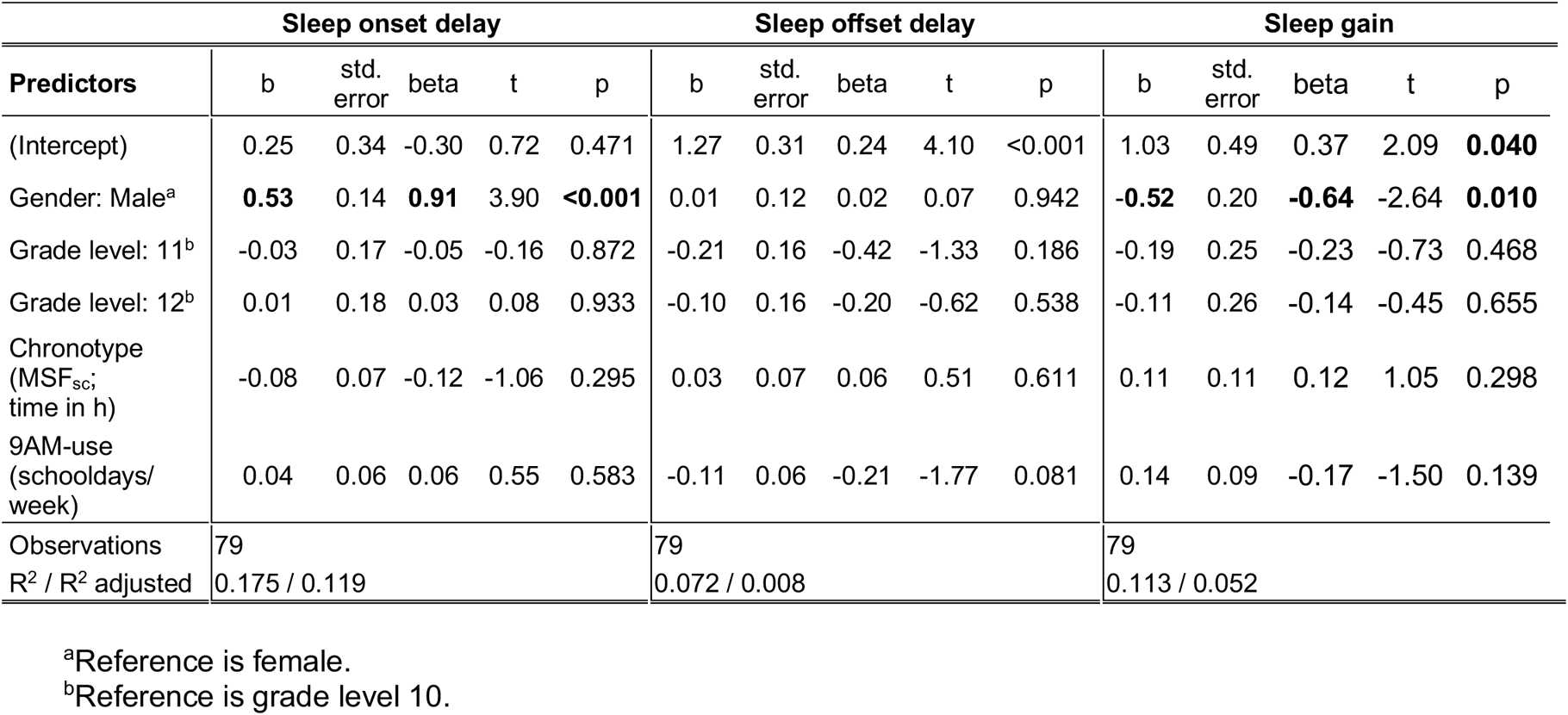
Individual differences in sleep gain on ≥9AM-days. Linear regression analyses on sleep gain, sleep onset delay and sleep offset delay on ≥9AM-days compared to 8AM-days in cohort 2 (N=79). Abbreviations: b, unstandardized coefficient; std. error, standard error; beta, standardized coefficient; t, t-statistic; p, p-value. R^2^ describes the explanatory power of the model (how much variance is explained). R^2^ adjusted is the explanatory power accounted for the number of predictors in the model.

| Predictors | Sleep onset delay |  |  |  |  | Sleep offset delay |  |  |  |  | Sleep gain |  |  |  |  |
| --- | --- | --- | --- | --- | --- | --- | --- | --- | --- | --- | --- | --- | --- | --- | --- |
|  | b | std. error | beta | t | p | b | std. error | beta | t | p | b | std. error | beta | t | p |
| (Intercept) | 0.25 | 0.34 | -0.30 | 0.72 | 0.471 | 1.27 | 0.31 | 0.24 | 4.10 | <0.001 | 1.03 | 0.49 | 0.37 | 2.09 | **0.040** |
| Gender: Male<sup>a</sup> | **0.53** | 0.14 | **0.91** | 3.90 | **<0.001** | 0.01 | 0.12 | 0.02 | 0.07 | 0.942 | **-0.52** | 0.20 | **-0.64** | -2.64 | **0.010** |
| Grade level: 11<sup>b</sup> | -0.03 | 0.17 | -0.05 | -0.16 | 0.872 | -0.21 | 0.16 | -0.42 | -1.33 | 0.186 | -0.19 | 0.25 | -0.23 | -0.73 | 0.468 |
| Grade level: 12<sup>b</sup> | 0.01 | 0.18 | 0.03 | 0.08 | 0.933 | -0.10 | 0.16 | -0.20 | -0.62 | 0.538 | -0.11 | 0.26 | -0.14 | -0.45 | 0.655 |
| Chronotype (MSF<sub>SC</sub>; time in h) | -0.08 | 0.07 | -0.12 | -1.06 | 0.295 | 0.03 | 0.07 | 0.06 | 0.51 | 0.611 | 0.11 | 0.11 | 0.12 | 1.05 | 0.298 |
| 9AM-use (schooldays/ week) | 0.04 | 0.06 | 0.06 | 0.55 | 0.583 | -0.11 | 0.06 | -0.21 | -1.77 | 0.081 | 0.14 | 0.09 | -0.17 | -1.50 | 0.139 |
| Observations | 79 |  |  |  |  | 79 |  |  |  |  | 79 |  |  |  |  |
| R<sup>2</sup> / R<sup>2</sup> adjusted | 0.175 / 0.119 |  |  |  |  | 0.072 / 0.008 |  |  |  |  | 0.113 / 0.052 |  |  |  |  |

<sup>a</sup>Reference is female.
<sup>b</sup>Reference is grade level 10.

**Table S3.** Sleep differences between time points and type of day in the longitudinal cohort (post-hoc comparisons relating to Fig. 6A). Two-way repeated measures ANOVAs were run for sleep onset, sleep offset, and sleep duration with the within-factors day (schooldays/weekends) and time point (t0/t1/t2) (see Fig. 6A). In case of significant interaction of both factors, simple effects (followed by Bonferroni-adjusted paired t-tests where indicated) are provided instead of the main effects. Data presented are mean ± standard deviation from the longitudinal cohort (n=33). η^2^, (partial) eta squared; d, Cohen’s d.

| Sleep onset |  |  |  |  |  |  |
| --- | --- | --- | --- | --- | --- | --- |
|  | t0 | t1 | t2 | Main effect:<br>*Day* | Simple effects: | Paired<br>t-tests: |
| Schooldays | -0.54h ± 0.79 | -0.43h ± 0.75 | -0.39h ± 0.73 | - | F(2,31)=1.61<br>p=0.217<br>η²=0.094, d=0.6442 | - |
| Weekends | 0.74h ± 0.94 | 0.46h ± 0.91 | 0.81h ± 1.00 |  | F(2,31)=2.68<br>p=0.084<br>η²=0.147, d=0.8303 | - |
| Main effect:<br>*Time point* | - |  |  | Main interaction<br>*Day\*Time point*:<br>F(2,64)=0.42<br>**p=0.020**<br>η²=0.013, d=0.230 |  |  |
| Simple effects: | F(1,32)=153.70<br>p<0.001<br>η²=0.823, d=4.313 | F(1,32)=62.57<br>p<0.001<br>η²=0.662, d=2.799 | F(1,32)=72.88<br>p<0.00<br>η²=0.695, d=3.019 |  |  |  |
| Sleep offset |  |  |  |  |  |  |
|  | t0 | t1 | t2 | Main effect:<br>*Day* | Simple effects: | Paired<br>t-tests: |
| Schooldays | 6.62h ± 0.45 | 6.80h ± 0.47 | 6.76h ± 0.44 | - | F(2,31)=9.03<br>**p=0.001**<br>**η²=0.368**, d=1.526 | t0-t1:<br>p<0.001<br>t0-t2:<br>p=0.025 |
| Weekends | 9.69h ± 0.97 | 9.31h ± 0.87 | 9.42h ± 0.93 |  | F(2,31)=4.88<br>**p=0.014**<br>**η²=0.240**, d=1.124 | t0-t1:<br>p=0.004 |
| Main effect:<br>*Time point* | - |  |  | Main interaction<br>*Day\*Time point*:<br>F(2,64)=10.42<br>**p<0.001**<br>**η²=0.246**, d=1.142 |  |  |
| Simple effects: | F(1,32)=294.21<br>**p<0.001**<br>**η²=0.902**, d=6.068 | F(1,32)=240.44<br>**p<0.001**<br>**η²=0.883**, d=5.494 | F(1,32)=322.85<br>**p<0.001**<br>**η²=0.910**, d=6.360 |  |  |  |
| Sleep duration |  |  |  |  |  |  |
|  | t0 | t1 | t2 | Main effect:<br>*Day* | Simple effects: | Paired<br>t-tests: |
| Schooldays | 7.10h ± 1.00 | 7.14h ± 0.56 | 7.16h ± 0.47 | - | F(2,31)=0.54<br>p=0.588<br>η²=0.034, d=0.375 | - |
| Weekends | 8.57h ± 0.42 | 8.50h ± 0.43 | 8.36h ± 0.55 |  | F(2,31)=2.70<br>p=0.083<br>η²=0.148, d=0.8336 | - |
| Main effect:<br>*Time point* | - |  |  | Main Interaction<br>*Day\*Time point*:<br>F(2,64)=3.88<br>**p=0.026**<br>**η²=0.108**, d=0.696 |  |  |
| Simple effects: | F(1,32)=120.24<br>**p<0.001**<br>**η²=0.790**, d=3.879 | F(1,32)=96.52<br>**p<0.001**<br>**η²=0.751**, d=3.473 | F(1,32)=59.07<br>**p<0.001**<br>**η²=0.649**, d=2.720 |  |  |  |

**Table S4.** Sleep differences between schooldays and weekends in cohort 2. Sleep onset, offset and duration from cohort 2 (n=105) at t2 are presented as mean ± standard deviation and were analysed via paired t-test. d_z_, Cohen’s d for paired t-tests.

|  | **Sleep onset** | **Sleep offset** | **Sleep duration** |
| --- | --- | --- | --- |
| **Schooldays** | -0.38h ± 0.81 | 6.83h ± 0.55 | 7.21 ± 0.76 |
| **Weekends** | 0.85h ± 1.11 | 9.57 ± 1.13 | 8.72h ± 0.95 |
| **Paired t-test** | t(104)=-14.757,<br>p<0.001<br>d<sub>z</sub>=1.440 | t(104)=-26.471,<br>p<0.001<br>d<sub>z</sub>=2.583 | t(104)=-14.230,<br>p<0.001<br>d<sub>z</sub>=1.389 |

## References

1. Crowley SJ, Van Reen E, LeBourgeois MK, et al. A longitudinal assessment of sleep timing, circadian phase, and phase angle of entrainment across human adolescence. PLoS One. 2014;9(11). doi:10.1371/journal.pone.0112199

2. Carskadon MA, Acebo C, Richardson GS, Tate BA, Seifer R. An Approach to Studying Circadian Rhythms of Adolescent Humans. J Biol Rhythms. 1997;12(3):278–289.

3. Carskadon MA, Acebo C, Jenni OG. Regulation of adolescent sleep: Implications for behavior. Ann N Y Acad Sci. 2004;1021(1):276–291. doi:10.1196/annals.1308.032

4. Jenni OG, Achermann P, Carskadon MA. Homeostatic sleep regulation in adolescents. Sleep. 2005;28(11):1446–1454. doi:10.1093/sleep/28.11.1446

5. Taylor DJ, Jenni OG, Acebo C, Carskadon MA. Sleep tendency during extended wakefulness: Insights into adolescent sleep regulation and behavior. J Sleep Res. 2005;14(3):239–244. doi:10.1111/j.1365-2869.2005.00467.x

6. Van Den Bulck J. Television viewing, computer game playing, and internet use and self-reported time to bed and time out of bed in secondary-school children. Sleep. 2004;27(1):101–104. doi:10.1093/sleep/27.1.101

7. Munezawa T, Kaneita Y, Osaki Y, et al. The Association between Use of Mobile Phones after Lights Out and Sleep Disturbances among Japanese Adolescents: A Nationwide Cross-Sectional Survey. Sleep. 2011;34(8):1013–1020. doi:10.5665/SLEEP.1152

8. Cajochen C. Alerting effects of light. Sleep Med Rev. 2007. doi:10.1016/j.smrv.2007.07.009

9. Souman JL, Tinga AM, te Pas SF, van Ee R, Vlaskamp BNS. Acute alerting effects of light: A systematic literature review. Behav Brain Res. 2018. doi:10.1016/j.bbr.2017.09.016

10. Yang M, Ma N, Zhu Y, et al. The acute effects of intermittent light exposure in the evening on alertness and subsequent sleep architecture. Int J Environ Res Public Health. 2018. doi:10.3390/ijerph15030524

11. Wittmann M, Dinich J, Merrow M, Roenneberg T. Social Jetlag: Misalignment of Biological and Social Time. Chronobiol Int. 2006;23(1-2):497–509. doi:10.1080/07420520500545979

12. Kuula L, Pesonen AK, Merikanto I, et al. Development of Late Circadian Preference: Sleep Timing From Childhood to Late Adolescence. J Pediatr. 2018;194:182–189.e1. doi:10.1016/j.jpeds.2017.10.068

13. Short MA, Gradisar M, Lack LC, et al. A Cross-Cultural Comparison of Sleep Duration Between U.S. and Australian Adolescents: The Effect of School Start Time, Parent-Set Bedtimes, and Extracurricular Load. Heal Educ Behav. 2013;40(3):323–330. doi:10.1177/1090198112451266

14. Hirshkowitz M, Whiton K, Albert SM, et al. National sleep foundation’s sleep time duration recommendations: Methodology and results summary. Sleep Heal. 2015;1(1):40–43. doi:10.1016/j.sleh.2014.12.010

15. Garaulet M, Ortega FB, Ruiz JR, et al. Short sleep duration is associated with increased obesity markers in European adolescents: Effect of physical activity and dietary habits. the HELENA study. Int J Obes. 2011. doi:10.1038/ijo.2011.149

16. Mullington JM, Haack M, Toth M, Serrador JM, Meier-Ewert HK. Cardiovascular, Inflammatory, and Metabolic Consequences of Sleep Deprivation. Prog Cardiovasc Dis. 2009. doi:10.1016/j.pcad.2008.10.003

17. Raniti MB, Allen NB, Schwartz O, et al. Sleep Duration and Sleep Quality: Associations With Depressive Symptoms Across Adolescence. Behav Sleep Med. 2017. doi:10.1080/15402002.2015.1120198

18. Short MA, Gradisar M, Lack LC, Wright HR. The impact of sleep on adolescent depressed mood, alertness and academic performance. J Adolesc. 2013. doi:10.1016/j.adolescence.2013.08.007

19. Baum KT, Desai A, Field J, Miller LE, Rausch J, Beebe DW. Sleep restriction worsens mood and emotion regulation in adolescents. J Child Psychol Psychiatry Allied Discip. 2014. doi:10.1111/jcpp.12125

20. Tynjälä J, Kannas L, Levälahti E. Perceived tiredness among adolescents and its association with sleep habits and use of psychoactive substances. J Sleep Res. 1997;6(3):189–198. doi:10.1046/j.1365-2869.1997.00048.x

21. Pasch KE, Latimer LA, Cance JD, Moe SG, Lytle LA. Longitudinal Bi-directional Relationships Between Sleep and Youth Substance Use. J Youth Adolesc. 2012. doi:10.1007/s10964-012-9784-5

22. Larcher S, Gauchez AS, Lablanche S, Pépin JL, Benhamou PY, Borel AL. Impact of sleep behavior on glycemic control in type 1 diabetes: The role of social jetlag. Eur J Endocrinol. 2016;175(5):411–419. doi:10.1530/EJE-16-0188

23. Parsons MJ, Moffitt TE, Gregory AM, et al. Social jetlag, obesity and metabolic disorder: Investigation in a cohort study. Int J Obes. 2015;39(5):842–848. doi:10.1038/ijo.2014.201

24. Levandovski R, Dantas G, Fernandes LC, et al. Depression Scores Associate With Chronotype and Social Jetlag in a Rural Population. Chronobiol Int. 2011;28(9):771–778. doi:10.3109/07420528.2011.602445

25. Wittmann M, Paulus M, Roenneberg T. Decreased Psychological Well-Being in Late ‘Chronotypes’ Is Mediated by Smoking and Alcohol Consumption. Subst Use Misuse. 2010;45(1-2):15–30. doi:10.3109/10826080903498952

26. Wahlstrom KL, Berger AT, Widome R. Relationships between school start time, sleep duration, and adolescent behaviors. Sleep Heal. 2017;3(3):216–221. doi:10.1016/j.sleh.2017.03.002

27. Lo JC, Lee SM, Lee XK, et al. Sustained benefits of delaying school start time on adolescent sleep and well-being. Sleep. 2018;(July):1–8. doi:10.1093/sleep/zsy052

28. Owens J. Insufficient Sleep in Adolescents and Young Adults: An Update on Causes and Consequences. Pediatrics. 2014;134(3):e921–e932. doi:10.1542/peds.2014-1696

29. Wheaton AG, Chapman DP, Croft JB, Chief B, Branch S. School start times, sleep, behavioral, health and academic outcomes: a review of literature. J Sch Heal. 2017;86(5):363–381. doi:10.1111/josh.12388.School

30. Bowers JM, Moyer A. Effects of school start time on students’ sleep duration, daytime sleepiness, and attendance: a meta-analysis. Sleep Heal. 2017;3(6):423–431. doi:10.1016/j.sleh.2017.08.004

31. Boergers J, Gable CJ, Owens JA. Later school start time is associated with improved sleep and daytime functioning in adolescents. J Dev Behav Pediatr. 2014. doi:10.1097/DBP.0000000000000018

32. Minges KE, Redeker NS. Delayed school start times and adolescent sleep: A systematic review of the experimental evidence. Sleep Med Rev. 2016;28:82–91. doi:10.1016/j.smrv.2015.06.002

33. Marx R, Tanner-Smith EE, Davison CM, et al. Later school start times for supporting the education, health, and well-being of high school students. Cochrane Database Syst Rev. 2017;2017(7). doi:10.1002/14651858.CD009467.pub2

34. Troxel WM, Wolfson AR. The intersection between sleep science and policy: introduction to the special issue on school start times. Sleep Heal. 2017;3(6):419–422. doi:10.1016/j.sleh.2017.10.001

35. Levin KA. Study design III: Cross-sectional studies. Evid Based Dent. 2006;7(1):24–25. doi:10.1038/sj.ebd.6400375

36. Illingworth G, Sharman R, Jowett A, Harvey CJ, Foster RG, Espie CA. Challenges in implementing and assessing outcomes of school start time change in the UK: experience of the Oxford Teensleep study. Sleep Med. 2019;60:89–95. doi:10.1016/j.sleep.2018.10.021

37. Estevan I, Silva A, Vetter C, Tassino B. Short Sleep Duration and Extremely Delayed Chronotypes in Uruguayan Youth: The Role of School Start Times and Social Constraints. J Biol Rhythms. 2020:1–14. doi:10.1177/0748730420927601

38. Goldin AP, Sigman M, Braier G, Golombek DA, Leone MJ. Interplay of chronotype and school timing predicts school performance. Nat Hum Behav. 2020:43–47. doi:10.1038/s41562-020-0820-2

39. Wahlstrom K. Changing Times: Findings From the First Longitudinal Study of Later High School Start Times. NASSP Bull. 2002;86(633):3–21. doi:10.1177/019263650208663302

40. Thacher P V., Onyper S V. Longitudinal Outcomes of Start Time Delay on Sleep, Behavior, and Achievement in High School. Sleep. 2016;39(2):271–281. doi:10.5665/sleep.5426

41. Lo JC, Lee SM, Lee XK, et al. Sustained benefits of delaying school start time on adolescent sleep and well-being. Sleep. 2018. doi:10.1093/sleep/zsy052

42. Das-Friebel A, Gkiouleka A, Grob A, Lemola S. Effects of a 20 minutes delay in school start time on bed and wake up times, daytime tiredness, behavioral persistence, and positive attitude towards life in adolescents. Sleep Med. 2020;66:103–109. doi:10.1016/j.sleep.2019.07.025

43. Widome R, Berger AT, Iber C, et al. Association of Delaying School Start Time with Sleep Duration, Timing, and Quality among Adolescents. JAMA Pediatr. 2020;174(7):697–704. doi:10.1001/jamapediatrics.2020.0344

44. Lufi D, Tzischinsky O, Hadar S. Delaying school starting time by one hour: Some effects on attention levels in adolescents. J Clin Sleep Med. 2011;7(2):137–143.

45. Nahmod NG, Lee S, Master L, Chang AM, Hale L, Buxton OM. Later high school start times associated with longer actigraphic sleep duration in adolescents. Sleep. 2019;42(2):1–10. doi:10.1093/sleep/zsy212

46. Dunster GP, de la Iglesia L, Ben-Hamo M, et al. Sleepmore in Seattle: Later school start times are associated with more sleep and better performance in high school students. Sci Adv. 2018. doi:10.1126/sciadv.aau6200

47. Carskadon MA, Wolfson AR, Acebo C, Tzischinsky O, Seifer R. Adolescent sleep patterns, circadian timing, and sleepiness at a transition to early school days. Sleep. 1998;21(8):871–881.

48. Winnebeck EC, Vuori-Brodowski MT, Biller AM, et al. Later school start times in a flexible system improve teenage sleep. Sleep. 2020;43:1–17. doi:10.1093/sleep/zsz307

49. Der Deutsche Schulpreis. Published 2019. https://www.deutscher-schulpreis.de/preistraeger/gymnasium-der-stadt-alsdorf

50. Ghotbi N, Pilz LK, Winnebeck EC, et al. The µMCTQ: An Ultra-Short Version of the Munich ChronoType Questionnaire. J Biol Rhythms. 2020;35(1):98–110. doi:10.1177/0748730419886986

51. Wickham H, Chang W, Henry L, et al. *R package ggplot2: Elegant Graphics for Data Analysis*. Version 3.2.1. 2019. https://cran.r-project.org/package=ggplot2

52. Pinheiro J, Bates D, DebRoy S, Sarkar D. *R package nlme: Linear and Nonlinear Mixed Effects Models.* Version 3.1-145. 2020. https://cran.r-project.org/package=nlme

53. Bates D, Maechler M, Bolker B, Walker S. *R package lme4: Linear Mixed-Effects Models using ‘Eigen’ and S4.* Version 1.1-18-1. 2018. https://cran.r-project.org/package=lme4

54. Paruthi S, Brooks LJ, D’Ambrosio C, et al. Recommended amount of sleep for pediatric populations: A consensus statement of the American Academy of Sleep Medicine. J Clin Sleep Med. 2016;12(6):785–786. doi:10.5664/jcsm.5866

55. Owens JA, Belon K, Moss P. Impact of delaying school start time on adolescent sleep, mood, and behavior. Arch Pediatr Adolesc Med. 2010;164(7):608–614. doi:10.1001/archpediatrics.2010.96

56. Wolfson A, Tzischinsky O, Brown C, Darley C, Acebo C, Carskadon M. Sleep, behavior, and stress at the transition to senior high school. Sleep Res. 1995;24:115.

57. Bai S, Karan M, Gonzales NA, Fuligni AJ. A daily diary study of sleep chronotype among Mexican-origin adolescents and parents: Implications for adolescent behavioral health. Dev Psychopathol. 2020:1–10. doi:10.1017/S0954579419001780

58. Crowley SJ, Wolfson AR, Tarokh L, Carskadon MA. An Update on Adolescent Sleep: New Evidence Informing the Perfect Storm Model. J Adolesc. 2018:55–65. doi:10.1016/j.adolescence.2018.06.001

59. Chellappa SL, Steiner R, Oelhafen P, Cajochen C. Sex differences in light sensitivity impact on brightness perception, vigilant attention and sleep in humans. Sci Rep. 2017;7(1):1–9. doi:10.1038/s41598-017-13973-1

60. Chung SJ, An H, Suh S. What do people do before going to bed? a study of bedtime procrastination using time use surveys. Sleep. 2020;43(4):1–10. doi:10.1093/sleep/zsz267

61. Estevan I, Silva A, Vetter C, Tassino B. Short Sleep Duration and Extremely Delayed Chronotypes in Uruguayan Youth: The Role of School Start Times and Social Constraints. J Biol Rhythms. 2020:0748730420927601.

62. Goldin AP, Sigman M, Braier G, Golombek DA, Leone MJ. Interplay of chronotype and school timing predicts school performance. Nat Hum Behav. 2020;4(4):387–396. doi:10.1038/s41562-020-0820-2

63. Biller AM, Molenda C, Obster F, et al. Are flexible school start times associated with higher academic grades? A 4-year longitudinal study. Submitted.

64. Campanini MZ, Lopez-Garcia E, Rodríguez-Artalejo F, González AD, Andrade SM, Mesas AE. Agreement between sleep diary and actigraphy in a highly educated Brazilian population. Sleep Med. 2017;35:27–34. doi:10.1016/j.sleep.2017.04.004

65. Tremaine RB, Dorrian J, Blunden S. Subjective and objective sleep in children and adolescents: Measurement, age, and gender differences. Sleep Biol Rhythms. 2010;8(4):229–238. doi:10.1111/j.1479-8425.2010.00452.x

66. Statistisches Bundesamt. Verteilung der Schüler auf Gymnasien nach dem höchsten Bildungsabschluss der Eltern im Jahr 2018. *Statista*. Published 2019. Accessed May 24, 2020. https://de.statista.com/statistik/daten/studie/162247/umfrage/besuch-des-gymnasiums-nach-abschluss-der-eltern/

